# Protocol for mapping of the supplementary motor area using repetitive navigated transcranial magnetic stimulation

**DOI:** 10.1101/2023.01.23.525160

**Authors:** Melina Engelhardt, Giulia Kern, Jari Karhu, Thomas Picht

**Affiliations:** Charité - Universitätsmedizin, corporate member of Freie Universität Berlin and Humboldt-Universität zu Berlin, Department of Neurosurgery, Charitéplatz 1, 10117 Berlin, Germany; Charité – Universitätsmedizin, corporate member of Freie Universität Berlin and Humboldt-Universität zu Berlin, Einstein Center for Neurosciences, Charitéplatz 1, 10117 Berlin, Germany; Charité – Universitätsmedizin, corporate member of Freie Universität Berlin and Humboldt-Universität zu Berlin, International Graduate Program Medical Neurosciences, Charitéplatz 1, 10117 Berlin, Germany; Department of Physiology, University of Eastern Finland, Kuopio, Finland; Cluster of Excellence Matters of Activity. Image Space Material, Humboldt-Universität zu Berlin, Unter den Linden 6, 10099 Berlin, Germany

**Author notes:** **Correspondence:** Melina Engelhardt. **Declarations of interest:** none.

**Keywords:** rTMS, Supplementary motor area, Brain mapping, Preoperative planning

## Abstract

**Background:** Damage to the supplementary motor area (SMA) for example during surgery can lead to impairments of motor and language function. A detailed preoperative mapping of functional boarders of the SMA could therefore aid preoperative diagnostics in these patients.

**Objective:** The aim of this study was the development of a repetitive nTMS protocol for non-invasive functional mapping of the SMA while assuring effects are indeed caused by SMA stimulation rather than activation of M1.

**Methods:** To this purpose the SMA in the dominant hemisphere of twelve healthy subjects (28.2 ± 7.7 years, 6 females) was mapped using repetitive nTMS at 20 Hz (120% RMT), while subjects performed a finger tapping task. The location of induced errors was marked in each subject’s individual MRI. To further validate the protocol, effects of SMA stimulation were directly compared to effects of M1 stimulation in four different tasks.

**Results:** Mapping of the SMA was possible for all subjects, yet varying effect sizes were observed. Stimulation of the SMA led to a significant reduction of finger taps compared to baseline (BL: 45 taps, SMA: 35.5 taps; p < 0.01). Line tracing, writing and targeting of circles was less accurate during SMA compared to M1 stimulation.

**Conclusion:** Mapping of the SMA using repetitive nTMS is feasible. While errors induced in the SMA are not entirely independent of M1 due to the proximity of both regions, disruption of the SMA induces functionally distinct errors. These error maps can aid preoperative diagnostics in patients with SMA related lesions.

## Introduction

Lesions of the supplementary motor area (SMA) can lead to widespread impairments such as unilateral motor deficits up to hemiplegia and mutism [1–3]. This so-called SMA syndrome is often temporary with patients recovering over the course of days to months. Yet, some patients report persisting deficits in complex motor functions several months after the lesion [2,4,5]. It has been suggested that functional reorganisation of the SMA mainly via recruitment of the contralateral SMA facilitates recovery of deficits [4,6–8]. However, these mechanisms are still poorly understood, and prediction of the individual extent and time course of recovery is limited.

Similarly, in case of surgical lesions to the SMA, preoperative assessment of the risk of inducing a SMA syndrome is limited. Determining the exact location of functionally relevant portions of the SMA can be crucial to avoid damaging these regions during surgery, specifically since the SMA is not limited by exact anatomical boundaries. Previous studies [9–11] used functional MRI to localize the exact location of the SMA on the individual patient’s brain. Yet these activation patters are often too large or unspecific to be used for surgical planning. Recently, repetitive TMS (rTMS) has been used to disrupt SMA function in healthy subjects [12,13]. While the spatial resolution of the reported protocol was still limited, it provided evidence that the SMA can be functionally mapped similarly to language and motor relevant areas [14,15]. Further, while single-neuron responses are similar between SMA and the primary motor cortex (M1), both areas show distinct population dynamics during execution of motor tasks [16]. Thus, disturbance of these dynamics using rTMS should lead to differential effects on task performance that can be used to separate SMA and M1 effects despite the vicinity of both areas.

The purpose of the present study was to develop a nTMS-based protocol to localize portions of the SMA relevant for motor function on the individual brain. Such a protocol could then be used in preoperative planning to preserve functional SMA areas, to assess the risk for postoperative SMA syndrome or to quantify the extent of postoperative reorganisation. Due to the proximity of SMA and M1 a specific focus of this study was to validate that induced effects are indeed caused by SMA stimulation rather than activation of M1.

## Material and methods

### Subjects

Twelve subjects (mean age 28.2 years, SD 7.7 years, 6 females) without any history of neurological or psychiatric illness provided their written informed consent to participate in this study. All subjects met the criteria for receiving an MRI scan and the TMS assessment. Exclusion criteria were history of epilepsy (also within the family), migraine, tinnitus, pregnancy, intake of prescription drugs within the past 14 days, permanent make-up, tattoos or metallic implants including any form of intrauterine devices. The study was conducted in accordance with the Declaration of Helsinki and approved by the local ethics committee.

### MRI

All subjects received a T1-weighted MPRAGE sequence (TR = 2.530 ms, TE = 4.94 ms, TI = 1.100 ms, flip angle = 7, voxel size = 1 mm × 1 mm × 1 mm, 176 slices) measured on a Siemens 3-T Magnetom Trio MRI scanner (Siemens AG, Erlangen, Germany). The scan took approximately 10 minutes for each subject.

### Neuronavigated TMS

The neuronavigated TMS (nTMS) assessment was divided into three parts: First the primary motor cortex was examined using single-pulse TMS. Next, the SMA was stimulated with repetitive TMS (rTMS) while subjects performed a motor task. In the final part, effects of stimulation of both areas were compared using different motor tasks. NTMS was applied using a Nexstim NBS 5 stimulator (Nexstim, Helsinki, Finland) with a figure-of-eight coil (outer diameter: 70 mm). The previously acquired structural MRI was used as a subject-specific navigational dataset.

### Motor assessment

Motor evoked potentials were recorded from the first dorsal interosseous muscle of the dominant hand. To this purpose, disposable Ag/AgCl surface electrodes (Neuroline 700; Ambu, Ballerup, Denmark) were attached in a belly-tendon fashion with the ground electrode on the left palmar wrist. Subjects were instructed to relax their hand muscles and muscle activity was monitored to assure relaxation of the muscle below a threshold of 10 µV. The motor hotspot was defined as stimulation site, electric field direction and angulation consistently eliciting the largest motor evoked potentials in the target muscle. For this point, the RMT was measured using the system’s inbuilt automated threshold hunting method [17]. The RMT was recorded as percentage of the stimulator output as well as the intensity of the induced electric field. To determine the size of the cortical representation of the target muscle, an area mapping with an intensity of 105% of the RMT was performed concentrically [18].

### SMA mapping

Consequently, anatomical boarders of the SMA region were estimated based on the structural MRI [19]. SMA was estimated as portion of the superior frontal gyrus until the point where a vertical line traversing the anterior commissure crosses the cortex. The suspected SMA region was then stimulated with rTMS (20 Hz, 120% RMT, 5s bursts, ITI 5s) while subjects performed a finger tapping task. We chose 20Hz as stimulation frequency as this was tolerated well and induced reliable disruptions of task performance in pilot subjects. For the finger tapping, subjects were instructed to tap with their dominant index finger as fast as possible. Each subject performed two rounds of finger tapping without stimulation as baseline. Next, the anatomically estimated SMA region was stimulated with rTMS. After 5 stimulations, subjects rested their hand for roughly one minute to prevent fatigue of the target muscles. Due to anatomical differences and different stages of protocol development, the amount of stimulation points varied between subjects. While we recommend around 20 stimulation points per hemisphere with each point being stimulated twice to assess replicability of the induced effects, SMA maps with fewer points are present in this study. The location of induced errors was marked in each subject’s individual MRI. Each session was further recorded on video to allow for offline analysis of induced errors using the nTMS systems inbuilt camera. The number of finger taps was recorded for each trial and converted to measure the reduction in finger taps compared to the baseline (in %).

### Comparison between SMA and M1 stimulation

To exclude SMA effects due to stimulation of M1 via the peripheral magnetic field, effects of SMA stimulation were directly compared to effects of M1 stimulation. To this purpose, a SMA hotspot was defined as the point eliciting the largest disruptions of task performance upon stimulation. For this point, the intensity of the induced electric field was recorded. Further, the electric field induced at the M1 hotspot when stimulating the SMA hotspot was recorded (residual SMA intensity). Consequently, the SMA hotspot was stimulated with the SMA mapping intensity (120% RMT) and the M1 hotspot was stimulated with the residual SMA intensity. Each subject performed two rounds without stimulation (baseline), with SMA stimulation and with M1 stimulation while executing 1 of 4 different tasks. Duration of stimulation was increased to last for the whole duration of the task. The order of the stimulation conditions and tasks applied were randomized between subjects.

Task 1 consisted of a finger tapping for 10 seconds. The number of finger taps was recorded as well as any noticeable deviations in the movement pattern. For task 2, subjects had to write a short sentence on a piece of paper. For analysis, the time to write the sentence and deviations in the writing pattern (legible, non-legible) were documented. An example categorization is presented in Fig. 3B. Task 3 required tracing a curved line with a pencil as fast and accurately as possible (Fig. S1). The task was stopped if subjects didn’t reach the end of the line after 20 seconds. Deviations from the line were analysed qualitatively (line traceable without problems, line traceable with strong deviations, line not traceable). An example categorization is presented in Fig. 3A. For the fourth task, subjects had to point a pencil to small circles on a paper as fast and accurately as possible for 20 seconds (Fig. S2). The number of circles targeted and number of circles missed (i.e. pencil marks outside a circle) were recorded.

**Fig. 1.**
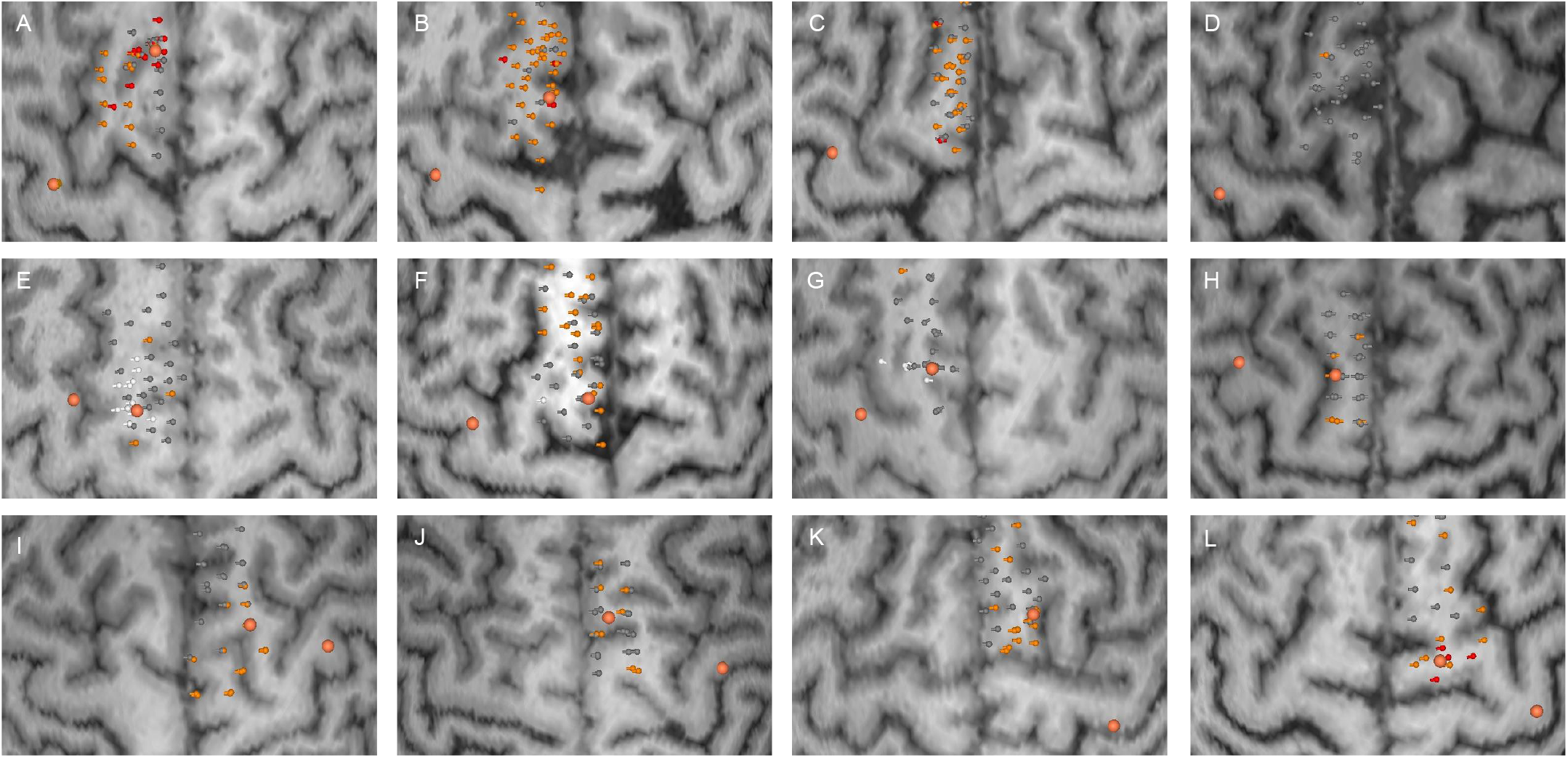
SMA error maps for all subjects. Each subplot (A-L) corresponds to one subject. Stimulation points are visualized with their respective error category in grey (no error), orange (mild errors) and red (significant errors). Stimulation points inducing an electric field above the RMT at M1 are marked in white and were excluded from SMA areas. Larger orange dots correspond to SMA and M1 hotspots respectively.

**Fig. 2.**
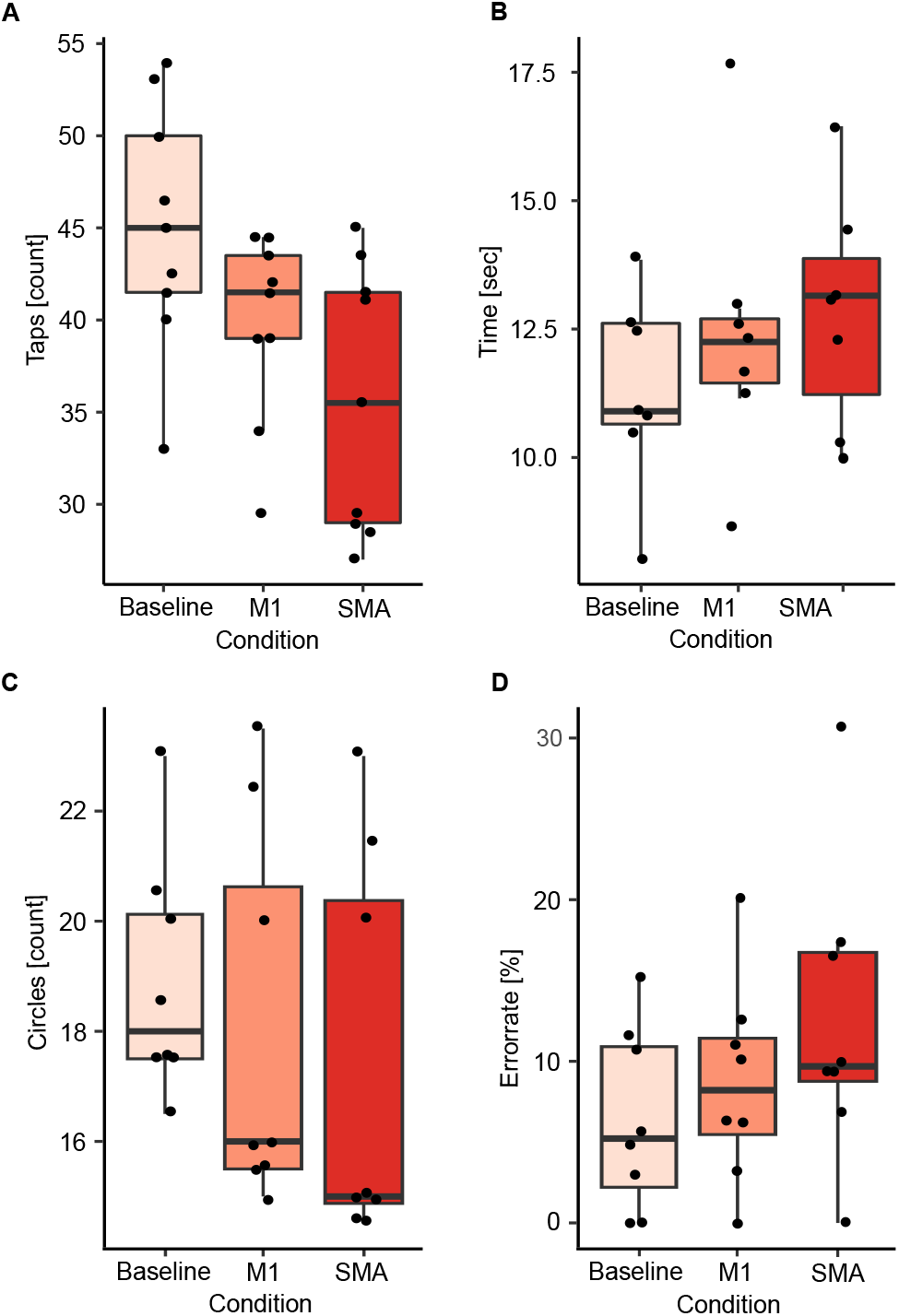
Comparison of SMA and M1 stimulation. Results are presented for the finger tapping task (A), the writing task (B), the number of circles targeted (C) and the error rate during the circle targeting task (D). Black dots correspond to average values for single subjects in each condition and task.

**Fig. 3.**
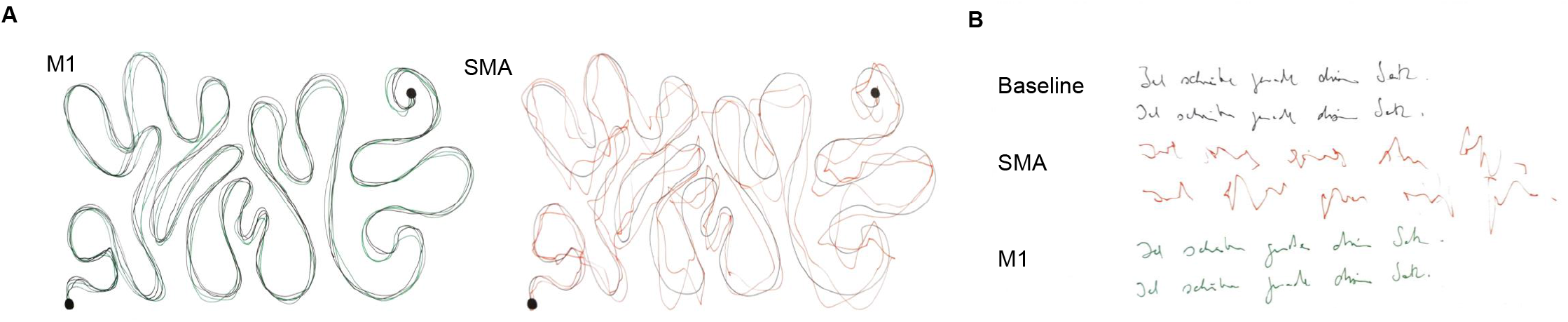
Example results of the line tracing (A) and writing task (B) for one subject. (A) The black line corresponds to the reference line during the tracing task. M1 stimulation was rated as traceable without problems and SMA stimulation as traceable with strong deviations. (B) Both baseline and M1 stimulation were rated as legible, whereas SMA stimulation was categorized as non-legible.

### Data analysis

The reduction in finger taps compared to baseline for each stimulation was categorized into three groups. Reductions ≤ 15% were counted as no errors, between 15% and 30% as mild errors, and > 30% as significant errors. These errors were marked using the nTMS system inbuilt analysis program and imported into the NBS software to create coloured SMA error maps.

Locations of SMA and M1 areas were assessed for potential overlaps in all subjects. We further compared the individual RMTs with the electric field induced at the M1 hotspot during SMA stimulation for all stimulation points. SMA stimulation points inducing an electric field above the RMT at M1 were highlighted and removed from SMA error maps.

For comparison of M1 and SMA stimulation, we first calculated the mean of the two trials per subject, task and stimulation condition. Next, values of baseline, M1 and SMA stimulation were compared using one-sided Wilcoxon signed-rank tests. Median values for each stimulation condition and task are reported. P-values < 0.05 were considered significant, p-values ≤ 0.1 are additionally reported as tendencies due to the small sample size. Further, qualitative deviations between both stimulations were analysed using the recorded videos and handwritten tests. These results are presented as incidence rates.

## Results

### SMA error maps

Errors in task performance could be induced in all 12 subjects. However, there was considerable variability in the size and location of SMA error maps as well as intensity of induced errors between subjects. Across subjects, a median of 34 points were stimulated (minimum 24, maximum 42 points). In 5 subjects, significant errors > 30% reductions could be induced over a median of 3 stimulation points. Accordingly, error incidence for significant errors varied between 3% and 26% between subjects with a median incidence of 7%. Mild errors (15 – 30% reduction) were recorded in all 12 subjects over a median of 11 stimulation points. Error incidence for mild errors varied between 3% and 79% between subjects with a median incidence of 36%. In two subjects (Fig. 1D, G), only one error could be induced after exclusion of errors likely due to M1 stimulation. SMA error maps of all subjects are presented in Figure 1.

### Comparison of SMA error maps and M1 areas

There was no overlap of stimulation points in SMA and M1 areas. However, in three subjects the electric field recoded at the motor hotspot was above the RMT for some of the functional points of the SMA area. In one subject, this included most of the functional SMA points as well as the SMA hotspot (Fig. 1E). The subject was therefore removed from further analysis. For the other two subjects, these points were highlighted and removed from the SMA error maps (Fig. 1F, G).

### Comparison of SMA and M1 stimulation

Definition of an SMA hotspot was possible in 9 out of 12 subjects. One subject had to be excluded as the electric field recorded at the motor hotspot was above the RMT for this location (Fig. 1E). In two other subjects, no clear hotspot could be detected during the SMA mapping (Fig. 1C, D). Finally, one of the remaining 9 subjects did not perform the writing task due to time constraints of the measurement.

A significant reduction of finger taps compared to baseline during SMA (BL: 45 taps, SMA: 35.5 taps; p < 0.01) and M1 stimulation (M1: 41.5 taps; p = 0.02) was observed. This effect was stronger during SMA then M1 stimulation (p = 0.04; Fig. 2A). Further, there was a tendency for a reduction in the number of circles targeted during SMA stimulation compared to baseline (BL: 18 circles, SMA: 15 circles; p = 0.09; Fig. 2C) as well as an increase in the error rate (BL: 5.2%, SMA: 9.7%; p = 0.07; Fig. 2D). No effects were observed during M1 stimulation for either the number of circles (M1: 16 circles; p = 0.24; Fig. 2C) or the error rate (M1: 8.2%; p = 0.32; Fig. 2D). Fewer circles were targeted during SMA compared to M1 stimulation (p < 0.01) and a tendency for a higher error rate was observed (p = 0.1). No differences between stimulations were observed in the time to complete the writing task (Fig. 2B).

Line tracing and writing was less accurate during SMA compared to M1 stimulation. In the line tracing task, 4 subjects (44%) were able to trace the line without deviations during stimulation of the SMA. 4 subjects (44%) showed strong deviations from the reference line and one subject (11%) was not able to trace the line at all. In contrast, during M1 stimulation only one subject showed slight deviations (11%) from the line. In the writing task, SMA stimulation led to a non-legible result in 3 subjects (33%), whereas only one subject produced non-legible writing during M1 stimulation. The error categories as well as results of both tasks for one example subject are presented in Fig. 3. In two subjects (40%) with visible disruptions in the line tracing task and one subject (33%) with disruptions in the writing task, effects of SMA disruption increased with stimulation time, while effects of M1 stimulation were present from the beginning on.

## Discussion

The present study developed a nTMS-based SMA mapping protocol with a high spatial resolution. In this way, we were able to localize functionally relevant subregions of the SMA within a larger anatomically predefined area. It was further possible to define a hotspot where the strongest errors could be induced, analogous to motor nTMS assessments.

The proposed protocol follows a virtual lesion paradigm as it is commonly used for the assessment of language function using nTMS [14,20]. Previously, two studies [12,13] in healthy subjects have shown that the SMA is susceptible to this kind of stimulation. In these studies, 10Hz stimulation could disrupt performance of different sub modalities of the Jebsen Taylor Hand function test when applied to the SMA. While these results are promising regarding the capabilities of rnTMS, stimulation was only applied to six predefined targets of the SMA. Therefore, it remained unclear if this stimulation paradigm can be extended to a detailed mapping of the SMA and whether it is sufficient to aid a presurgical planning in case of lesions affecting the SMA region. To address this question, the present study used a short 5-second finger tapping task which can be repeated over multiple stimulation points to assess SMA function rather than a set of multiple tasks. In favour of this task, the SMA has been suggested to play a role in the encoding of movement sequences [1,21] and fMRI studies have shown an activation of the SMA during finger tapping [9]. Further, rTMS has been used to modulate inter-tap intervals during finger tapping [13]. While the present study focused on the number of taps as mapping outcome, further studies should certainly look into a more detailed analysis of movement kinematics to make mapping more specific to different functional aspects.

Using this stimulation protocol, it is possible to achieve a high-resolution mapping in roughly 10 minutes for one hemisphere. The proposed protocol includes short breaks after a maximum of five stimulation targets to avoid fatigue of hand muscles. Further, to improve accuracy of the functional assessment, each stimulation target should be stimulated at least twice. Classification of errors could then be restricted to points with replicable reductions in finger tapping performance, thus increasing reliability of the assessment. Future studies should also investigate the proposed limits for error categories to quantify when a reduction in finger taps is sufficiently large. This could be aided by studies in neurosurgical patients, where resection of functional points can be compared with occurrence and severity of functional impairments.

In the present study, a disruption of performance was also visible in other tasks involving more complex movements such as writing or targeting circles. An SMA mapping using these tasks would require more time. Yet our findings highlight the possibility to study somatotopy and functional organisation of the SMA using rnTMS, when paired for example with toe tapping to study lower extremity function or targeting small circles to examine more complex coordinated movements. Previous studies have suggested that the SMA has a somatotopic organisation with lower extremities being represented in the posterior SMA, upper extremities in the medial SMA and the face in the most anterior part of the SMA [11]. While the present study was not designed to investigate SMA somatotopy, most subjects showed the strongest disruptions in task performance in medial to posterior portions of the SMA thus supporting this notion. Since additional tasks in this study were only tested on the SMA hotspot, no further effects of somatotopy could be investigated. However, these effects should be studied more systematically by including lower extremity movements to assess any spatial separation of effects. Additionally, it could be investigated whether any hemispheric differences are present in the functional organisation of the SMA.

Importantly, there was a considerable variation in the strengths of the induced stimulation effect between subjects. We hypothesize that in some subjects the applied stimulation intensity might not have been sufficient to disrupt larger portions of the SMA region. As the SMA is located in the posterior portion of the superior frontal gyrus extending into the interhemispheric cleft [1,19], some subregions might be more difficult to stimulate as they are further away from the coil. In the present study, we refrained from increasing the stimulation intensity beyond 120% of the RMT due to the novelty of the protocol, to reduce the risk for any side effects and to reduce the risk of M1 contamination of the effects. However, these modifications might have been necessary to induce stronger effects in some subjects. Since the stimulation was tolerated well in our study, it seems that these limits can be exceeded in future studies and thereby the responder rate might be increased if effects are controlled for M1 contamination.

Finally, we aimed to distinguish effects of SMA and M1 stimulation to ensure recorded stimulation effects are not due to an indirect activation of the primary motor cortex [22]. In support of our protocol, errors induced during stimulation of the SMA hotspot were stronger than during M1 stimulation with the residual SMA targeting electric field. Further, at least in some subjects effects induced by SMA stimulation were qualitatively different from M1 stimulation. Errors built up over time, while effects of M1 stimulation were present from the beginning on. However, an induced electric field larger than the RMT was recorded over the motor hotspot when stimulating some of the functionally positive points over the SMA. This is not surprising given the proximity of both regions and intensity of stimulation but warrants caution when interpreting any SMA mapping results. Consequently, we argue that the electric field induced at M1 should always be controlled for when stimulating the SMA. In this way, validity of the SMA mapping can be ensured. Future studies in patients could further assess whether stimulated points are functionally essential by comparing resection of positive nTMS mapping points to occurrence and type of postoperative deficits [23]. Once this relationship is established, nTMS based SMA mapping could be integrated in preoperative planning and risk stratification as well as to assess postoperative reorganisation and rehabilitation of deficits.

## Conclusions

In conclusion, mapping of the supplementary motor area using high frequency rnTMS is possible as stimulation of the SMA can disrupt hand movements analogous to a virtual lesion paradigm. Due to the proximity to the primary motor cortex, stimulation intensities as well as the electric field induced at M1 need to be monitored during SMA mapping so assure validity of the induced errors. Finally, this protocol could also be integrated in preoperative planning to address the functional relevance of TMS-positive SMA parts, to assess the risk for developing a postoperative SMA syndrome or to quantify the extent of postoperative reorganisation.

## CRediT author statement

Melina Engelhardt: Conceptualization, Methodology, Formal analysis, Investigation, Data Curation, Writing – Original Draft, Visualization, Project administration. Giulia Kern: Formal analysis, Data Curation, Writing – Review & Editing. Jari Karhu: Conceptualization, Methodology, Writing – Review & Editing. Thomas Picht: Conceptualization, Methodology, Writing – Review & Editing, Supervision.

## Acknowledgements

MR-imaging for this study was performed at the Berlin Center for Advanced Neuroimaging (BCAN).

