## Supplementary Materials for "Protocol for mapping of the supplementary motor area using repetitive navigated transcranial magnetic stimulation"

*** Correspondence:**

Melina Engelhardt

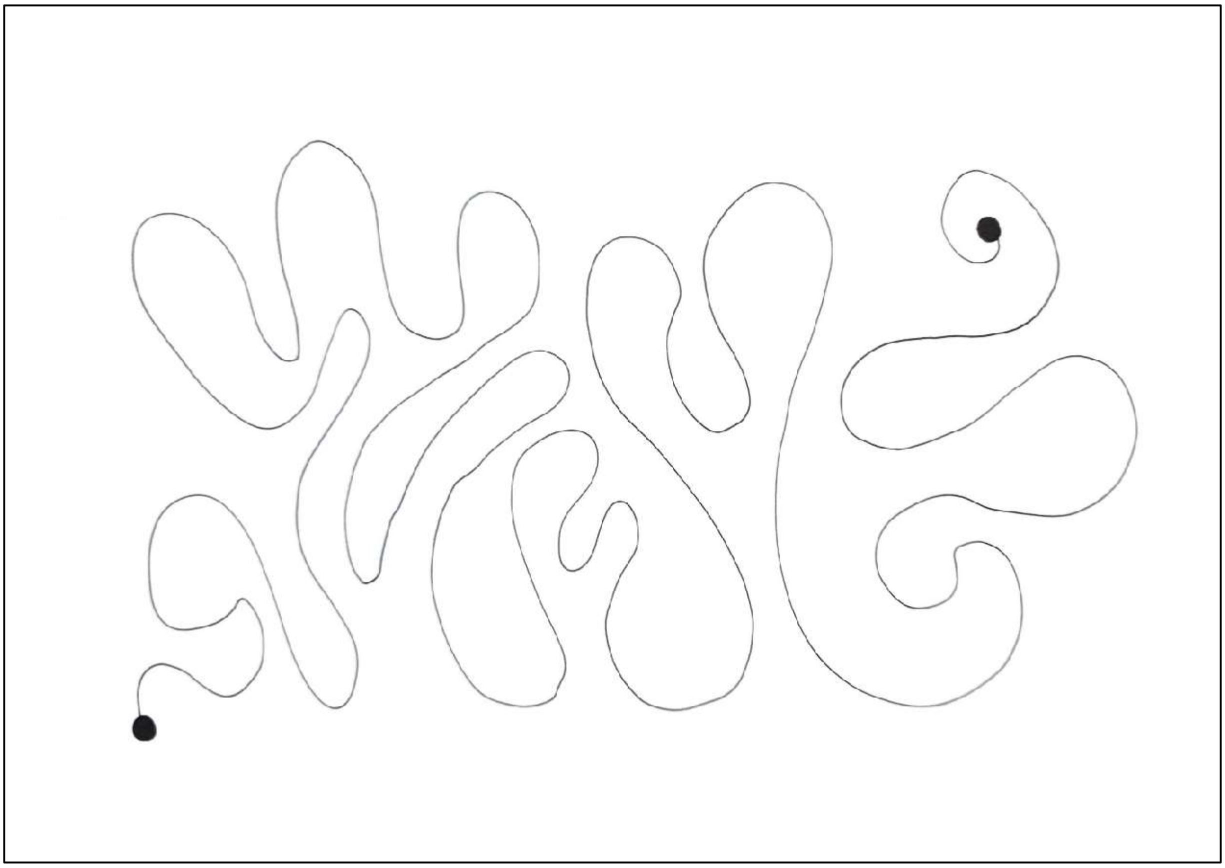

**Fig. S1** Task template for line tracing. Subjects were instructed to trace the line as fast and accurately as possible starting in the bottom left corner. Stimulation was applied for 20 seconds or until the end of the line was reached.

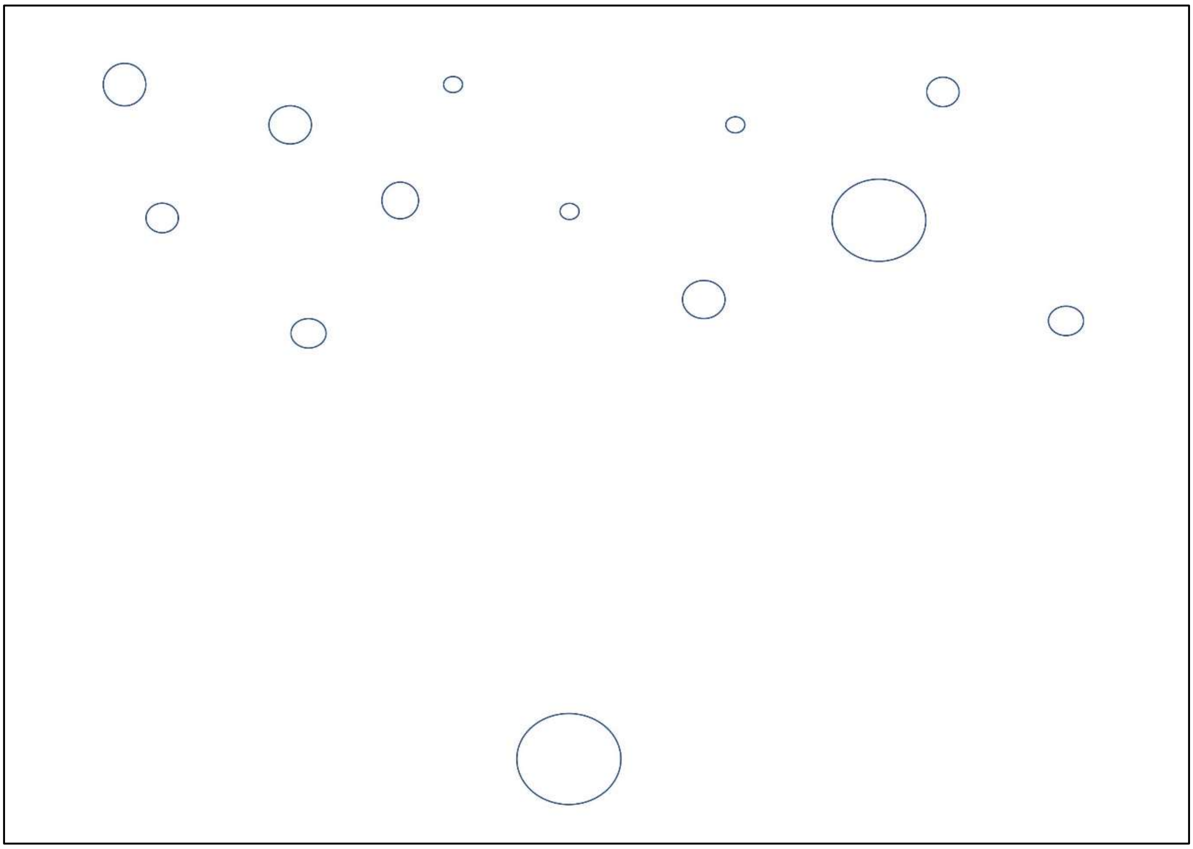

**Fig. S2** Task template for circle targeting. Subjects were instructed to mark the circles with a pencil starting with the big circle on the bottom of the page. They then had to alternate between the larger bottom circle and the circles on the top of the page until all circles were marked. Stimulation was applied for 20 seconds.
